# Inferring viral proteins that act as public goods during coinfection

**DOI:** 10.64898/2026.07.02.736036

**Authors:** Yael Maoz, Moran Meir, Nadav Ben Nun, Yoav Ram, Adi Stern

## Abstract

Interactions among individuals in structured populations can alter fitness effects of mutations and reshape evolutionary processes. In many systems, including bacteria, yeast, and viruses, such interactions often result in public goods: gene products that are costly to produce yet exploitable by others. During viral coinfection of the same cell, gene products from one genome may complement deleterious mutations in another, allowing defective genomes to persist. Yet it remains difficult to infer which proteins are shareable from population sequencing data, because mutation, selection, drift, and complementation are intertwined. Here, we developed a quantitative framework to infer protein-specific public goods in the RNA bacteriophage MS2, which encodes only four proteins. We analyzed experimental evolution data generated under two multiplicity-of-infection (MOI) regimes: low MOI, where coinfection is rare, and high MOI, where coinfection is common. We first compared empirical mutation patterns between regimes and then applied a Wright-Fisher model combined with simulation-based Bayesian inference using neural posterior estimation. In a two-stage strategy, gene-specific fitness effects were inferred from low-MOI data and subsequently used to estimate protein sharing under high-MOI conditions. Across two statistical inference frameworks, lysis emerged as the strongest public-good candidate, replicase and coat showed an intermediate signal, and maturation showed the weakest evidence for sharing. Together, our results show that viral proteins differ markedly in their propensity to act as public goods. More broadly, they illustrate how coinfection can generate density-dependent selection, a general feature of social evolution that may shape evolutionary dynamics.

## Introduction

In structured populations, interactions among individuals, including co-infecting genomes, can alter the fitness effects of mutations and reshape evolutionary processes. Such interactions arise whenever individuals share a local environment, allowing gene products produced by one individual to affect others. Viruses provide a particularly tractable example of this principle. During coinfection, multiple viral genomes infect the same cell, coexist in a shared intracellular environment, and can interact through the products they encode. These interactions can alter the fitness effects of mutations and, in turn, change the course of viral evolution and ecology (Díaz-Muñoz, 2017; Leeks et al., 2023; Sanjuán, 2021; Sanjuán et al., 2004). One important class of such interactions is complementation. During complementation, a genome carrying a deleterious or even lethal mutation is rescued by a coinfecting genome that supplies the missing functional product in trans (Segredo-Otero & Sanjuán, 2022) (Fig. 1). In evolutionary game-theory terms, the defective genome exploits a resource that it did not produce, and the relevant gene product can therefore be viewed as a public good. Recent work has further emphasized that the evolutionary consequences of coinfection depend on which viral products are shared among genomes within the same cell (Leeks et al., 2021; Robertson et al., 2026). At the extreme end of this process lie defective interfering particles and other social cheaters, which not only persist in mixed infection but can outcompete wild-type genomes by avoiding the costs of producing shared functions or by gaining additional intrinsic advantages (Leeks et al., 2021; Meir et al., 2020).

**Figure 1.**
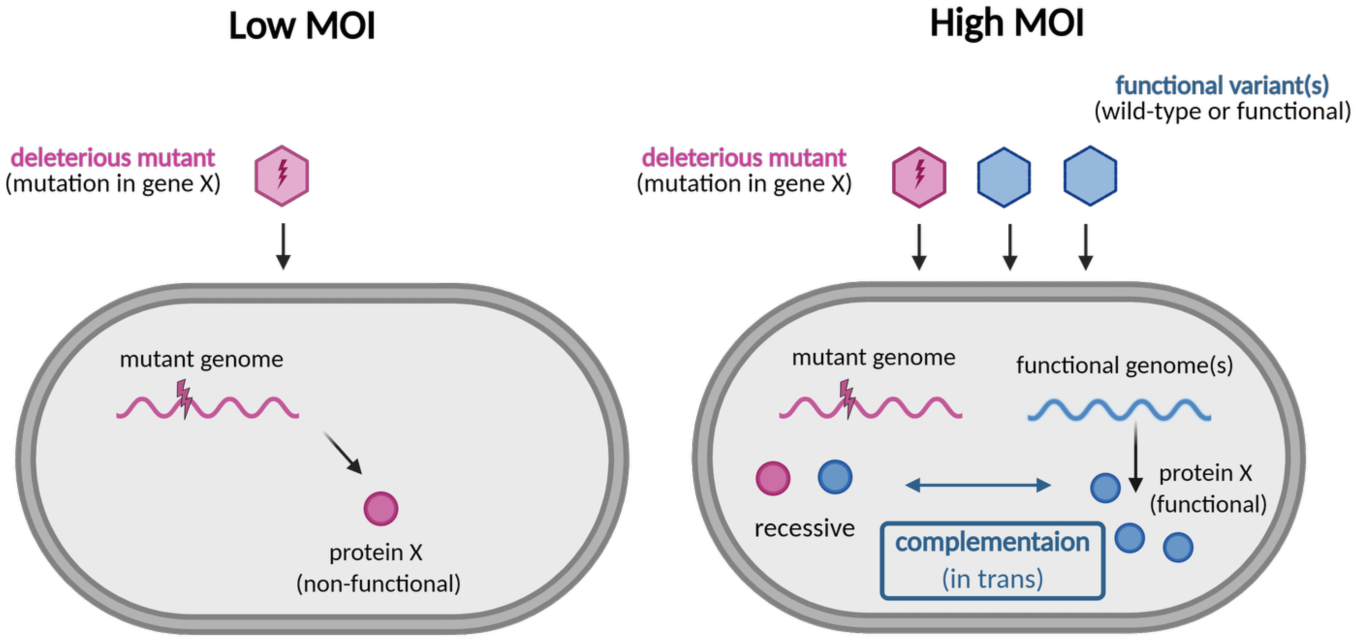
Conceptual model for complementation during viral coinfection. At low MOI, deleterious mutant genomes are typically exposed to selection because they infect cells without a complementing functional genome. At high MOI, coinfecting functional genomes can supply the missing protein product in trans, masking the deleterious effect of the mutation and allowing it to behave as recessive. The extent of this masking is expected to depend on the viral protein affected by the mutation.

Not all viral functions are expected to be equally shareable. Some proteins may act largely in cis and remain tightly associated with the genome that encoded them. Others may diffuse or operate at the level of the infected cell and thereby become accessible to all genomes present. Distinguishing between these cases is important because shareable functions should experience relaxed purifying selection under frequent coinfection (Chao & Elena, 2017; Díaz-Muñoz et al., 2017). Deleterious mutations affecting such proteins may behave as effectively recessive: they reduce fitness in singly infected cells but have little or no effect when complemented by coinfecting wild-type genomes (Froissart et al., 2004; Gao & Feldman, 2009).

The extent to which these interactions matter depends strongly on multiplicity of infection (MOI), defined as the average number of infectious particles infecting a host cell. At low MOI, most infected cells receive a single genome and selection largely reflects the intrinsic fitness effects of mutations (Sanjuán et al., 2004). At high MOI, coinfection becomes common and genomes experience density-dependent selection, one in which complementation, exploitation, and interference can all occur (Koelle et al., 2019; Martin et al., 2020; Sanjuán, 2017; Turner & Chao, 1999). Contrasting low- and high-MOI evolution experiments therefore offers a controlled way to examine how intracellular social interactions reshape mutational fitness effects.

Here, we focused on the RNA bacteriophage MS2, a positive-sense single-stranded RNA virus with a compact 3,569-nt genome that encodes only four proteins: maturation, coat, lysis, and replicase (Fiers et al., 1976). MS2 is a particularly attractive system (Betancourt, 2009) because its molecular biology has been well studied, its compact genome allows full-length haplotype sequencing, and our previous studies have already shown that lysis-defective and replicase-defective cheaters can spread under high MOI (Meir et al., 2020, 2025).

MS2 infection begins when the maturation protein mediates attachment to the F pilus and entry into the host cell. Once inside, the genomic RNA serves directly as mRNA. The coat protein (also called cp) forms the capsid but also participates in regulating translation of the replicase gene, whereas the replicase protein is the viral catalytic subunit of the RNA-dependent RNA polymerase that copies the genome using host factors. The lysis protein is expressed late and triggers host-cell lysis, releasing all progeny virions. These four proteins therefore differ not only in function but also in timing, abundance, and likely spatial coupling to the genome that encoded them (Duin & Tsareva, 2005; Stockley et al., 2013; Valegård et al., 1990).

These differences make it unclear a priori which functions should be most shareable during coinfection. Lysis is the clearest candidate for a public good because it acts at the level of the whole infected cell (in *trans*), so once sufficient lysis protein is present, all progeny in that cell can benefit (Chamakura & Young, 2020; Meir et al., 2020; Young, 1992). For the other proteins, the balance between *cis* and *trans* action is less obvious. Replicase may act in *trans* because polymerase produced from one genome can in principle copy another genome, yet translation-replication coupling could create a local advantage for the parental template (Ahlquist, 2002). Coat protein is diffusible and requires 178 copies, which could favor sharing, but its roles in RNA recognition, translational regulation, and virion assembly may still generate genome-specific constraints (Valegård et al., 1990). Maturation protein is an especially ambiguous case: because it is produced at low stoichiometry and has a specialized role in entry and virion function, it may be difficult to exploit, yet some of its contributions to particle production could still be complemented indirectly in mixed infections. We therefore view protein sharing not as a binary property but as a continuum, ranging from largely *cis*-acting to highly *trans*-complementable functions.

In principle, such differences in shareability should leave a signature in population sequencing data, because complementation can relax the effective selective constraint on mutations in shared genes. However, inferring such protein-specific sharing from sequence data is not straightforward. Observed mutation frequencies are shaped simultaneously by mutation, selection, genetic drift (population size, passage bottlenecks), and sequencing noise (Gillespie, 2004). A statistical framework that explicitly models these processes is therefore needed.

Here, we combined an empirical statistical analysis of mutation frequencies with a mechanistic evolutionary model and simulation-based Bayesian inference to quantify the extent to which each MS2 protein behaves as a public good. We first analyzed low-frequency mutations in low- and high-MOI long-read sequencing data using a hierarchical beta-binomial framework to estimate protein-specific effects of coinfection on deleterious and synonymous mutations. We then developed a two-stage mechanistic evolutionary model and fit it using simulation-based Bayesian inference. In stage 1, baseline mutation and selection parameters were inferred from low-MOI data, where complementation is assumed negligible, and in stage 2, gene-specific complementation parameters were inferred from high-MOI data. Together, these complementary approaches allowed us to ask whether the four MS2 proteins differ in their propensity to function as intracellular public goods.

## Results

### Mutation patterns at high MOI suggest complementation in specific genes

To assess how coinfection shapes mutational fitness effects, we analyzed longitudinal experimental evolution data from MS2 populations propagated under low (MOI = 0.1) and high (MOI = 10) MOI conditions, using long-read sequencing that spans the entire genome at passage 10 from independently evolved lines. By this stage, we have previously shown that adaptive mutations rose to appreciable frequencies (typically >3%) (Meir et al., 2020, 2025), providing sufficient signal to infer selection from sequence data. Under low MOI, most cells are infected by a single genome, whereas high MOI promotes frequent coinfection and potential complementation.

We began by comparing mutation patterns between these regimes to test for signatures of relaxed purifying selection under coinfection. Because such signatures are expected to be most visible among rare variants, we focused on low-frequency mutations, defined as mutations below 1% but above the detection limit of approximately 0.1% (Methods). The rationale was that deleterious mutations should remain rare or undetectable under low MOI, but may accumulate into this measurable frequency range under high MOI if their effects are masked by complementation. Hundreds of low-frequency mutations were observed in both conditions, but their composition differed across genes and MOI regimes (Fig. S1). As an initial descriptive analysis, we examined line-level mean frequencies of low-frequency nonsynonymous mutations at passage 10, separately for each protein. These mutations tended to reach higher frequencies under MOI = 10 than under MOI = 0.1 in lysis, coat, and replicase, whereas maturation did not show a comparable increase (Fig. 2A). By contrast, synonymous mutations showed no consistent MOI-dependent differences across genes. This distinction is important because synonymous mutations should be less sensitive to functional complementation than deleterious nonsynonymous mutations.

**Figure 2.**
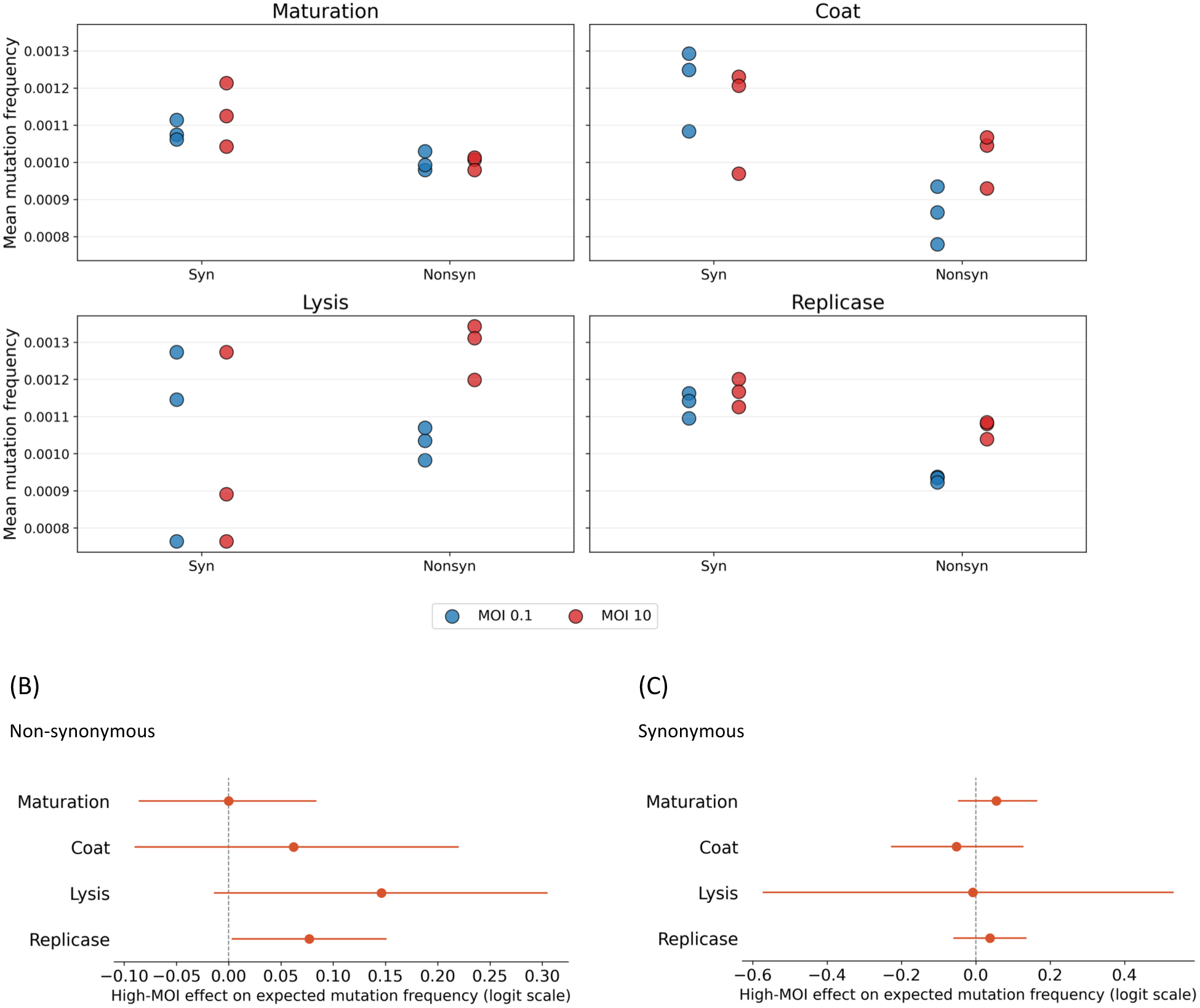
Evidence for altered mutational fitness effects under high MOI. (A) For each MS2 protein, points show the line-level mean frequency of passage-10 low-frequency mutations in independently evolved populations under MOI = 0.1 (blue) or MOI = 10 (red). Low-frequency mutations were defined as mutations below 1% frequency and above the detection threshold after down-sampling to equal coverage. Synonymous and nonsynonymous mutations are shown separately. (B, C) Estimates of the high-MOI effect on low-frequency nonsynonymous (B) and synonymous (C) mutations. Protein-specific high-MOI effects were estimated using a hierarchical beta-binomial model fitted to low-frequency, non-adaptive mutant read counts. Points indicate posterior means and horizontal bars indicate 95% highest density intervals (HDIs). The dashed vertical line marks no effect.

Because mutation-frequency data are bounded, sparse, and overdispersed, we next used a hierarchical beta-binomial model to formally evaluate these patterns. The model was fitted directly to mutant read counts rather than to line-level mean frequencies (Methods). For each low-frequency, non-adaptive nonsynonymous mutation, the model estimated the effect of high MOI separately for each protein, while accounting for overdispersion and including experimental line as a random effect. This analysis supported a protein-specific increase in nonsynonymous mutation frequencies under high MOI (Fig. 2B). Among nonsynonymous mutations, replicase showed a 95% HDI entirely above zero, and lysis showed a strongly positive effect whose 95% HDI only marginally overlapped zero. Coat also had a positive effect, but its 95% HDI overlapped zero more substantially. Finally, maturation was overall centered around zero. By contrast, none of the synonymous high-MOI effects had 95% HDIs excluding zero, and effects were concentrated around zero (Fig. 2C). Thus, high MOI was associated with increased frequencies of low-frequency nonsynonymous mutations in lysis and replicase, weak evidence for such an increase in coat, and no evidence for an increase in maturation.

### The empirical signal is robust to the mutation-frequency cutoff

Because the upper frequency threshold of 1% used to define low-frequency variants is somewhat arbitrary, we repeated the hierarchical beta-binomial analysis using two additional cutoffs: 0.5% and 2%. For nonsynonymous non-adaptive mutations, the qualitative pattern was stable across thresholds, high MOI had little or no effect in maturation, a positive but uncertain effect in coat, and stronger positive effects in lysis and replicase (Fig. S2A). In contrast, the synonymous-mutation analysis did not show a comparable protein-specific pattern consistent with complementation (Fig. S2B). Synonymous effects were generally centered near zero or had broad uncertainty intervals, and their direction was not consistently aligned with the nonsynonymous signal. Overall, these threshold analyses indicate that the main empirical conclusion does not depend on the exact frequency cutoff used to define low-frequency mutations. High MOI is associated primarily with increased frequencies of nonsynonymous mutations in lysis and replicase, with weaker support for coat and no clear effect in maturation.

### Mutations under high MOI do not preferentially target the most conserved sites

If complementation under high MOI mainly rescued strongly deleterious mutations, one might expect high-MOI mutations to be concentrated at especially conserved sites. We therefore compared conservation scores of nonsynonymous mutated sites between MOI regimes for each gene. Across all four proteins, we did not detect significant differences in the conservation levels of mutated sites between MOI = 10 and MOI = 0.1 (Table S1). Moreover, we did not observe a clear relationship between mutation frequencies at MOI = 0.1 and site-specific conservation, suggesting that conditions in our experimental system do not closely follow long-term evolutionary constraints inferred from natural sequence variation.

### A two-stage inference framework separates intrinsic fitness effects from complementation

Although the empirical analyses were consistent with relaxed purifying selection under coinfection, they also underscored the need for a mechanistic model. The beta-binomial results showed protein-specific positive high-MOI effects for nonsynonymous mutations, but with varying uncertainty across proteins. Direct comparisons of mutation frequencies therefore could not by themselves distinguish complementation from altered mutation input, or replicate-level variation. Furthermore, differences between MOI regimes could in principle arise from changes in mutation rate or in the number of replication cycles, rather than from complementation itself. Indeed, we observed a slight difference in the accumulation of synonymous mutations between conditions (Fig. S3). This raised the possibility that higher mutation frequencies under high MOI could be driven by altered mutation input (e.g., higher mutation rates) rather than relaxed selection.

To disentangle these causes, we developed an inference framework that explicitly links changes in genotype frequencies to selection and complementation under different MOI regimes. It builds on our previously developed framework for MS2 evolution under low-MOI conditions, in which mutation rates and selection parameters were inferred from haplotype sequencing data (Caspi et al., 2023). Here, we extend that framework to incorporate protein-specific complementation under co-infection. In this framework, we model a viral population evolving under serial passaging using a Wright-Fisher model in which individuals are defined by the number and class of mutations they carry. Rather than modeling whole-genome sequences, we represent each genotype by counts of mutant alleles classified by gene, type (synonymous vs nonsynonymous), and fitness effect (adaptive vs deleterious) (Methods). To describe how deleterious mutations behave under coinfection, we borrow the terms dominant and recessive from diploid genetics. Specifically, deleterious nonsynonymous mutations in the viral population are treated as dominant when their fitness effects remain exposed to selection despite coinfection, and as recessive when their effects can be masked by complementation.

In this framework, complementation is captured by allowing deleterious mutations in a given gene to behave as effectively neutral with a gene-specific probability under high MOI. This recessive probability provides a quantitative measure of how shareable each protein function is within coinfected cells. Evolution then proceeds through mutation, selection, and sampling across passages, generating expected distributions of mutation frequencies that can be compared directly to the empirical summary statistics. To infer model parameters from the data, we used simulation-based Bayesian inference (Cranmer et al., 2020) via neural posterior estimation (NPE). In this approach, the model is repeatedly simulated across a wide range of parameter values, and a neural density estimator is trained to approximate the posterior distribution of parameters given the observed summary statistics. This allows joint inference of mutation, selection, and complementation parameters while using an evolutionary model with an intractable likelihood function.

A key challenge is that the apparent relaxation of purifying selection under high MOI could reflect either masking due to complementation or differences in the underlying mutation accumulation process between regimes. To separate these effects, we implemented a two-stage inference strategy. In the first stage, the model is fitted to low-MOI data, where coinfection is rare, to infer baseline parameters, including the mutation rate and gene-specific fitness effects of deleterious nonsynonymous mutations (termed stage 1). In the second stage, these parameters are used to define informative (narrow) priors for the high-MOI inference, while the gene-specific deleterious fitness effects are held fixed at their low-MOI estimates (Methods). The model is then fitted to the high-MOI data to re-infer the mutation rate and to estimate gene-specific probabilities that deleterious mutations behave as recessive under coinfection (termed stage 2). This design allows differences between MOI regimes to be attributed specifically to complementation, while accounting for potential differences in mutation input and baseline selection across genes.

### Synthetic-data benchmarks support the accuracy of the inference framework

We first evaluated the inference framework on synthetic datasets generated under the corresponding low and high MOI models. Across both stages, 95% highest density intervals showed empirical coverages close to the nominal level, albeit with a mild tendency toward overconfidence. Maximum-a-posteriori (MAP) estimates were largely unbiased, with log10 (MAP/true) values centered near zero for most parameters (Table S2; Fig. S4). Small deviations were observed in inferred adaptive probabilities and adaptive fitness effects, which is expected because these parameters are partially correlated. Overall, the synthetic benchmarks indicated that the framework can estimate the true parameters with sufficient accuracy for biological interpretation.

### High-MOI inference reveals coinfection-specific sharing in lysis, coat, and replicase

We next applied the two-stage inference framework to the empirical data. In stage 1, we inferred mutation and selection parameters from the low-MOI data, where coinfection is expected to be rare, thereby estimating the intrinsic fitness effects of deleterious nonsynonymous mutations in each protein. In stage 2, we fitted the high-MOI data while holding these protein-specific deleterious fitness effects fixed at their low-MOI estimates, allowing us to infer gene-specific recessive probabilities under coinfection.

To test whether the inferred recessive probabilities captured a genuine high-MOI signal rather than an intrinsic tendency of the inference procedure to assign non-zero sharing, we performed a leave-one-replicate-out low-MOI control analysis. In this control, the same inference framework used for the high-MOI data was applied to low-MOI trajectories, where coinfection is rare and complementation is not expected. Specifically, for each low-MOI replicate in turn, the remaining low-MOI replicates were used to infer the baseline mutation and fitness parameters, and the withheld replicate was then treated as the target dataset for estimating an apparent recessive probability. This provides a matched null distribution: the degree of “sharing” inferred by the model when the target data come from low-MOI evolution and should therefore contain little true complementation signal.

The empirical high-MOI posteriors were shifted above the low-MOI control posteriors for all four proteins, but the magnitude of this shift differed substantially across proteins (Fig. 3, Table 1). Lysis showed by far the strongest separation: the high-MOI posterior was concentrated near very high recessive probabilities, whereas the low-MOI control remained close to zero, consistent with extensive masking of deleterious lysis mutations under coinfection. The remaining proteins all showed substantially weaker shifts. Replicase exhibited the clearest separation from its low-MOI control despite intermediate inferred recessive probabilities, coat showed a somewhat larger inferred effect, and maturation displayed the smallest shift. Together, this analysis provides strong evidence for extensive sharing of lysis, supports intermediate sharing of replicase and coat with differing degrees of confidence, and provides only limited evidence for sharing of maturation.

**Figure 3.**
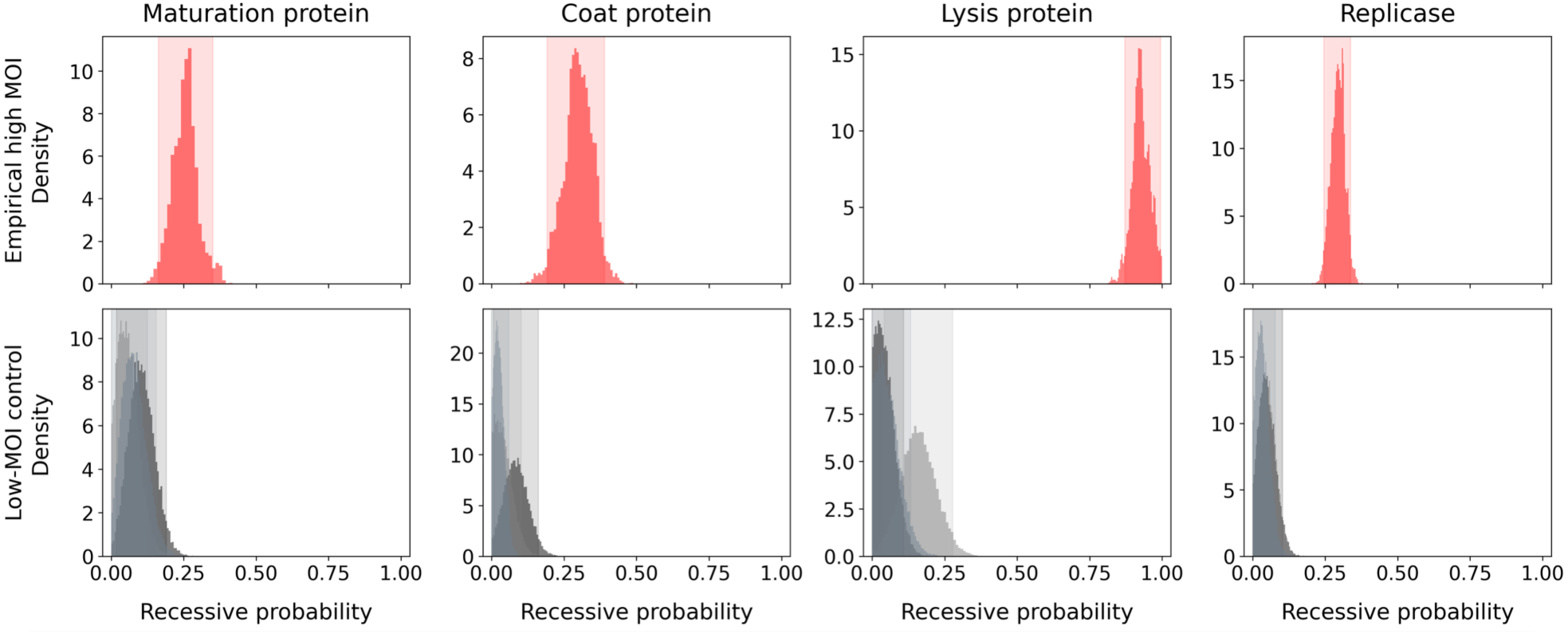
High-MOI inference of protein-specific recessive probabilities compared with low-MOI controls. Posterior distributions are shown for the probability that deleterious nonsynonymous mutations behave as recessive under coinfection for each MS2 protein. The upper row shows the empirical high-MOI posterior (in red) inferred from passage-10 MOI = 10 data using a collective posterior across the three high-MOI lines. The lower row shows low-MOI control posteriors (grey shade for each line) inferred in a leave-one-out design, in which the same high-MOI inference framework was applied to low-MOI data despite coinfection being rare. Boxed shaded regions indicate 95% highest density intervals.

**Table 1.** Inferred recessive probabilities for deleterious nonsynonymous mutations under high MOI and corresponding low-MOI control estimates. Values shown for the high-MOI analysis are posterior maximum-a-posteriori (MAP) estimates with 95% highest density intervals (HDIs). For the low-MOI control, the table reports the largest upper bound of the 95% HDI obtained across the leave-one-replicate-out control analyses for each protein.

|  | High MOI | Control upper HDI |
| --- | --- | --- |
| Maturation | 0.24 [0.163,0.35] | 0.190 |
| Coat | 0.345 [0.191,0.389] | 0.161 |
| Lysis | 0.915 [0.871,0.994] | 0.277 |
| Replicase | 0.313 [0.245,0.338] | 0.012 |

Beyond the public-goods parameters themselves, the high-MOI inference of the mutation rate (MAP of 0.269 mutations/genome/replication cycle) also yielded a slightly higher value than the low-MOI stage (MAP of 0.253), in line with the genomic data analysis (Fig. S2) and repeatedly assigned a relatively high adaptive-mutation probability to lysis.

Posterior predictive checks provided an additional validation step. Simulations performed under the inferred posterior distributions reproduced the observed mutation summary statistics for both low-MOI (Fig. S5) and high-MOI datasets with good fidelity (Fig. 4), indicating that the model captures the main statistical structure of the data even though it is necessarily a simplified representation of MS2 evolution.

**Figure 4.**
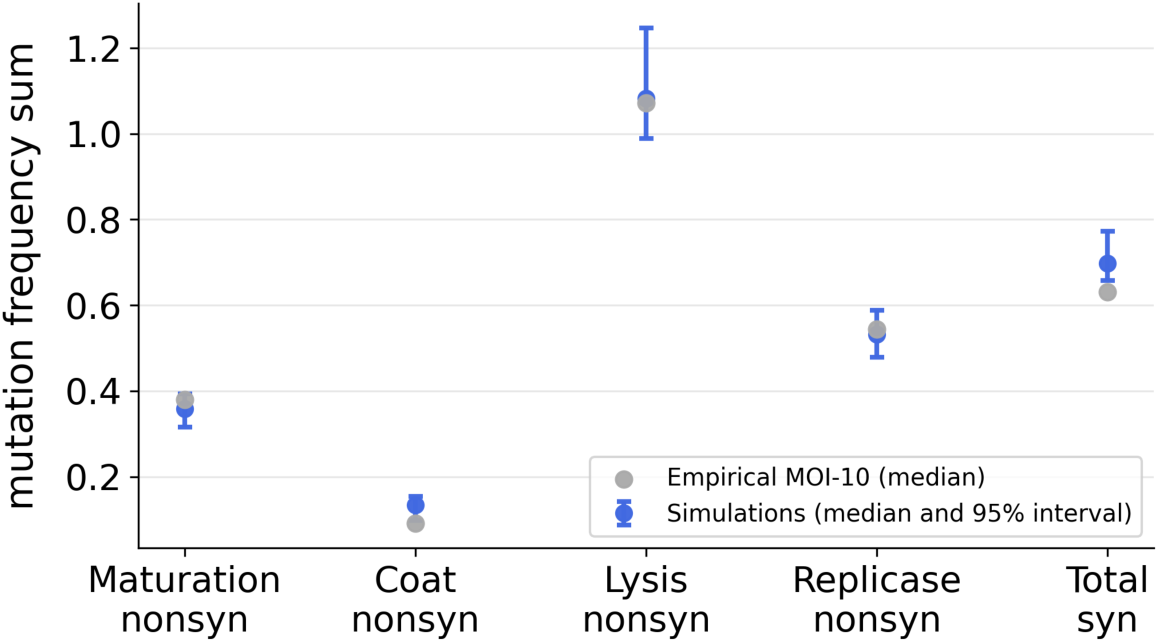
The inferred evolutionary model reproduces the observed mutation patterns. Comparison of empirical mutation frequencies sum at passage 10 (gray) to 200 simulations generated using parameters sampled from the inferred posterior distributions derived from the evolutionary model (blue) under MOI 10. Points show the median value for each mutation class, and blue error bars indicate the 95% simulation interval.

## Discussion

Our results support a simple but important conclusion: under frequent coinfection, viral proteins can behave as intracellular public goods, but they do so to markedly different extents. Together, the two complementary modeling frameworks support a hierarchy of protein sharing within MS2-infected cells. Lysis showed the strongest and most consistent evidence for extensive sharing and complementation, replicase exhibited consistent evidence for partial sharing, coat displayed an intermediate estimated effect accompanied by greater uncertainty, and maturation showed little evidence that it is shared. This pattern implies that coinfection can relax purifying selection in a strongly gene-specific manner.

The inference for lysis protein behaving as public goods is biologically intuitive. Lysis acts at the level of the infected cell, and once sufficient lysis protein accumulates, all progeny virions are released regardless of which genome encoded the functional lysis protein (Young, 1992). In that sense, lysis is a canonical public good (Meir et al., 2020, 2025). The enrichment of nonsense mutations in lysis under high MOI and the near-complete masking inferred by the model both fit this expectation. The result also argues against dominant-negative lysis mutants being common enough to overwhelm complementation, because such mutants would be expected to depress the inferred recessive probability.

Replicase and coat appear to occupy a more intermediate position. One useful way to think about this is as a continuum between locality and diffusibility. Replicase may often act locally near the genome from which it was translated (Ahlquist, 2002), which would limit sharing, yet our previous work has shown that replicase-defective cheaters can nonetheless be complemented under coinfection (Meir et al., 2020, 2025). Importantly, this complementation was incomplete: replication remained strongly limited even in coinfected cells and was only compensated by a substantial advantage in packaging. This suggests that replicase is only partially diffusible and can act in trans to a limited extent (Nagy & Simon, 1997; Simon-Loriere & Holmes, 2011), consistent with our inference of intermediate sharing. In contrast, coat presents a different type of constraint: because many copies (n=178) are required to build a virion (Valegård et al., 1990), complete exploitation is not guaranteed even if coat proteins diffuse freely. Partial sharing therefore seems plausible for both proteins, albeit for different mechanistic reasons.

The weakest signal for maturation is also informative. The maturation protein is produced at very low levels and its expression is tightly regulated by RNA structure (Duin & Tsareva, 2005; Stockley et al., 2013). A product that is scarce, temporally restricted, or tightly coupled to the genome that encodes it should be harder to exploit (Chao & Elena, 2017). Our results are consistent with that expectation. We do not interpret the maturation estimates as proving the complete absence of complementation, but rather as indicating that any such effect is much weaker than for the other genes and difficult to distinguish from zero with the available data.

An important strength of this study is the combination of two frameworks for statistical inference. The mutation count analysis alone already suggest relaxed purifying selection under high MOI, but could in principle be confounded by gene-specific differences in baseline fitness effects or by differences in mutation accumulation across conditions. The two-stage inference with the evolutionary model addresses this by anchoring intrinsic deleterious effects in the low-MOI regime and only then estimating coinfection-specific masking. The low-MOI leave-one-out control further strengthens the interpretation by showing that the model does not routinely infer strong recessiveness when coinfection is absent.

At the same time, several limitations should be kept in mind. Biological replicate variation was evident for both MOI=0.1 and MOI=10 data (Fig. 2A), and the MOI=10 experiment had long-read data only at passage 10, limiting temporal resolution. Our categorization of adaptive versus non-adaptive mutations is necessarily approximate, and the model makes simplifying assumptions, including shared fitness effects within mutation classes and an abstract genotype representation (Beaumont, 2010; Gillespie, 2004). These simplifications are what make inference feasible, but they also mean that the inferred recessive probability should be interpreted as an effective summary of complementation propensity rather than as a direct biochemical measurement.

The broader implication is that viral fitness landscapes are conditional on infection context. A mutation that is strongly deleterious in singly infected cells may behave nearly neutrally in coinfected cells if the relevant function is shareable. This context dependence should influence the maintenance of genetic diversity, the persistence of defective genomes, and the emergence of social cheaters. In natural systems, where multiplicity and spatial structure vary over time and across tissues, such effects may be especially important.

In summary, we provide strongest evidence that lysis functions as a public good during MS2 coinfection, together with intermediate evidence for replicase and coat, whereas maturation appears substantially less shareable. More generally, we introduce a framework for quantifying protein-specific complementation from viral evolution experiments. With suitable adaptations, the same strategy could be extended to other RNA viruses and to other forms of intracellular interaction, helping to bridge mechanistic virology with evolutionary theory on cooperation, cheating, and density-dependent selection.

## Methods

### Experimental evolution datasets

We analyzed previously generated MS2 serial passaging experiments performed under two infection regimes: MOI = 0.1 and MOI = 10 (Caspi et al., 2023; Meir et al., 2020, 2025). All lines were initiated from clonal MS2 stocks derived from single plaques to minimize standing variation at the start of the experiment. Phages were propagated on naïve Escherichia coli c-3000 hosts, with fresh host cultures used at each passage to minimize host adaptation and host-phage coevolution. Experiments were carried out for ten serial passages under controlled population sizes and controlled MOI.

For low MOI, we used passage-10 long-read data from three independently evolved lines. For high MOI, eight biological replicates were performed (replicas A-H), and four of these were sequenced at passage 10 (A, E, G, H) (Meir et al., 2025). To allow for balanced data we used only three of the passage 10 sequenced population (E, G, H).

### Long-read sequencing and mutation calling

RNA from evolved populations was sequenced using the LoopSeq synthetic long-read approach, which reconstructs full-length viral haplotypes using molecular barcodes and short-read sequencing (Callahan et al., 2021). Long reads were aligned to the MS2 reference genome using BLAST (Altschul et al., 1990; McGinnis & Madden, 2004) as described previously (Caspi et al., 2023). Only coding regions were included in downstream analyses, together accounting for more than 90% of the genome. Indels were excluded because their functional effects are more difficult to model in a reduced genotype framework. Founder mutations previously identified in the low-MOI dataset were also removed to avoid confounding the inference with standing genetic variation already present in the initiating stock.

Mutations were classified as synonymous or nonsynonymous, and repeatedly rising mutations (in two lines or more) that reached a frequency of 3% or higher were labelled as adaptive based on their frequencies across replicate lines (Caspi et al., 2023; Meir et al., 2025). To make lines directly comparable despite differences in sequencing depth, each line was down-sampled to the minimum coverage observed across the analyzed datasets (n = 1309 genomes). Mutations in overlapping reading frames were counted once and assigned to their nonsynonymous instance.

### Empirical analyses

We first performed non-model-based comparisons of mutation burdens between low and high MOI. Unless stated otherwise, these analyses were restricted to mutations with frequencies above ∼0.1% (or more precisely, 1/1309 since this was our limit of detection; see below) but below 1% at passage 10. These analyses were performed separately for nonsynonymous non-adaptive mutations and for synonymous mutations. Because mutation frequencies are bounded, sparse, and overdispersed, we used a hierarchical beta-binomial model rather than a t-test to compare mutation frequencies between MOI regimes while accounting for variation among experimental lines.

For each mutation *l* in experimental line *e*, let *y_e,l_* denote the number of mutant reads and *n_e,l_* (1309) the total number of reads covering that position. We modeled the mutant read counts as

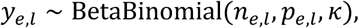

where *p_e,l_* is the expected mutation frequency and *κ* is an overdispersion parameter. The expected frequency was modeled on the logit scale as

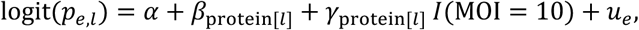

where *α* is the global intercept, *β*_protein[*l*]_ is the baseline effect for the protein in which mutation *l* occurs (maturation, coat, lysis, or replicase), *I*(MOI = 10) is an indicator variable equal to 1 for MOI = 10 and 0 for 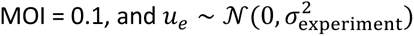 is a random intercept for each experimental line. The protein-specific high-MOI effects *γ*_protein[*l*]_, were the parameters of interest, because they quantify whether low-frequency mutation frequencies increase under coinfection for each protein. Posterior distributions of these effects were summarized using posterior means, 95% highest density intervals, and the posterior probability that the high-MOI effect was greater than zero.

To test whether high-MOI mutations occur preferentially at strongly constrained positions, we compared site-specific conservation scores of mutated nonsynonymous sites between MOI conditions. Conservation scores were taken from a previous conservation analysis of MS2 and related viruses (Meir et al., 2025), that was based on inferring site-specific evolutionary rates (Mayrose et al., 2004; Pupko et al., 2002).

### Evolutionary model

We developed a stochastic Wright-Fisher model of MS2 evolution during serial passaging (Fisher, 1923; Wright, 1931). Explicit nucleotide-level tracking of all genotypes is computationally infeasible, so genotypes were represented in a reduced but biologically motivated form (Beaumont, 2010; Gillespie, 2004). Each mutation was classified by mutational type (synonymous or nonsynonymous), by adaptive status, and, for deleterious nonsynonymous mutations, by the gene they affect and whether they behave as dominant or recessive under coinfection.

In the full model, each genotype is therefore represented by counts of gene-specific deleterious nonsynonymous recessive mutations, gene-specific deleterious nonsynonymous dominant mutations, gene-specific adaptive nonsynonymous mutations, and genome-wide counts of adaptive and non-adaptive synonymous mutations. Genotypes sharing the same mutation-count vector are treated as equivalent genotype classes (Fig. 5). This abstraction greatly reduces the state space while preserving the features required to infer differences among genes in effective complementation.

**Figure 5.**
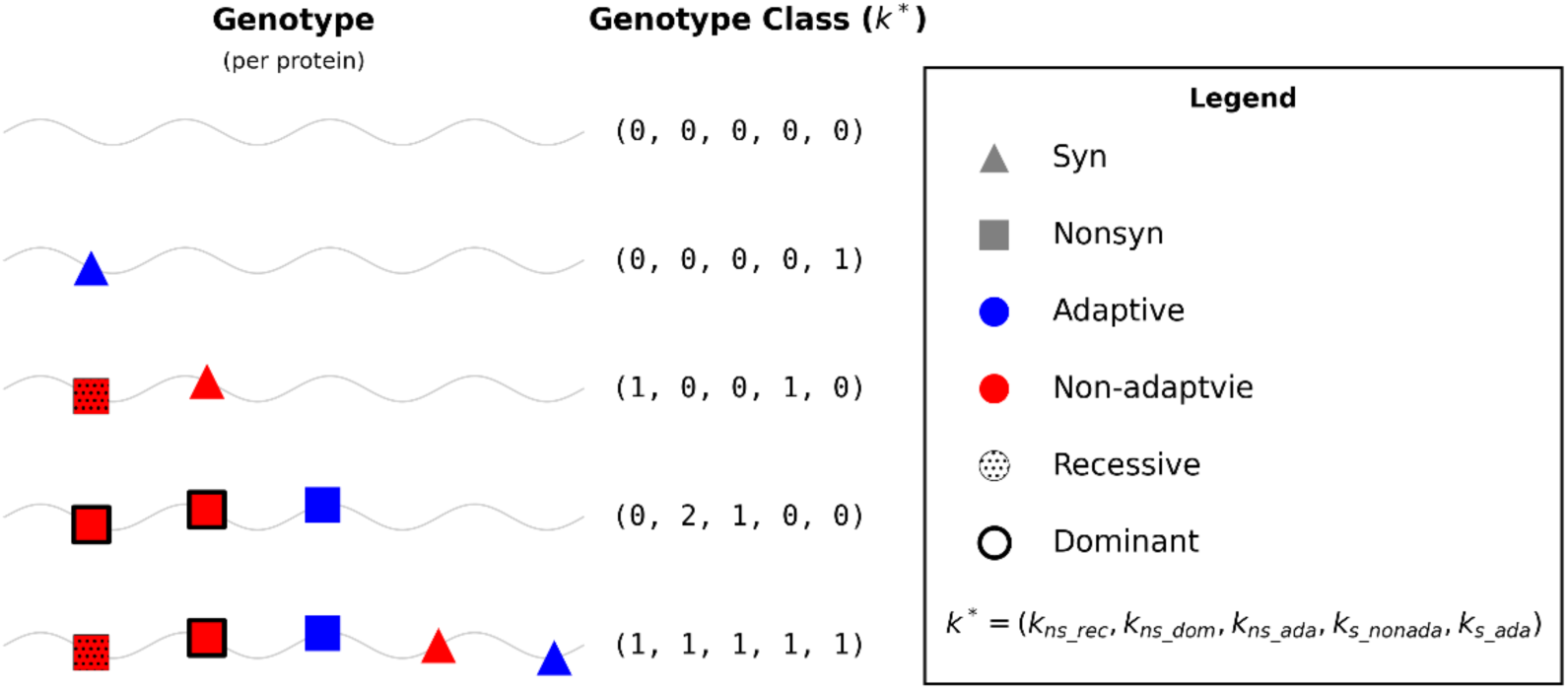
Illustration of simplified genotype classes. Genotypes in the model are grouped into genotype classes according to the types and counts of mutations they contain. In the full model, each genotype is represented by a 14-dimensional vector composed of: for each gene (maturation, coat, lysis, and replicase), the number of nonsynonymous non-adaptive recessive mutations, nonsynonymous non-adaptive dominant mutations, and nonsynonymous adaptive mutations; and two additional genome-wide quantities corresponding to the total number of synonymous non-adaptive and synonymous adaptive mutations. For clarity, the illustration depicts mutations in a single gene only. In the actual model, synonymous mutations are counted genome-wide rather than per gene. The simplified genotype class *k*^∗^is therefore represented as a 5-tuple corresponding to the counts of each mutation category under a single-gene abstraction. Within each category, SNVs are assumed to be interchangeable and to share the same fitness effect (*ω*). Grouping genotypes into classes substantially reduces the dimensionality of the state space, while preserving a detailed representation of the underlying evolutionary dynamics.

We model serial passaging as a Wright-Fisher process with constant population size N and discrete, non-overlapping generations. In each generation, genotype frequencies are updated through mutation, selection, and random sampling.

#### Mutation

The number of new mutations per genome per replication cycle is assumed to follow a Poisson distribution *u* ∼ *Poisson*(*U*) where U is the genome-wide mutation rate. Mutations are distributed across genes according to their coding lengths and are partitioned into mutation classes (synonymous/nonsynonymous, adaptive/non-adaptive, dominant/recessive) according to fixed probabilities. Mutation updates genotype-class frequencies by redistributing mass between classes according to these transition probabilities.

#### Fitness and complementation

Fitness is assumed to be multiplicative across mutation classes. For a genotype k, fitness is given by

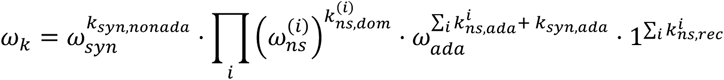

Here, *k* indexes a genotype class, defined by its mutation-count vector. The index *i* runs over genes (e.g., maturation, coat, lysis, replicase).

A key feature of the model is the treatment of deleterious nonsynonymous mutations under coinfection. Mutations classified as recessive are assumed to be fully complemented and therefore do not reduce fitness (i.e., contribute a factor of 1), whereas dominant mutations reduce fitness according to gene-specific effects. The probability that a deleterious nonsynonymous mutation is recessive is gene-specific and constitutes the main parameter capturing public-goods behavior.

#### Selection

Following mutation, genotype frequencies are updated deterministically according to selection:

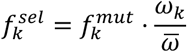

Where *f*_k_ is the frequency of genotype class *k* in the population before selection and 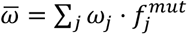 is the mean population fitness.

#### Genetic drift and serial passaging

Genetic drift and experimental bottlenecks are modeled by multinomial sampling of N genomes:

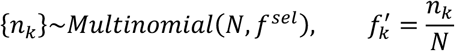

Here, *n_k_* is the number of genomes of genotype class *k* after sampling, and 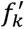 is the frequency of genotype class *k* in the next generation. This sampling step captures stochasticity arising from finite population size and the serial transfer process.

#### Sequencing process

To account for the fact that only a subset of genomes is observed and that sequencing is error-prone, we explicitly model the sequencing process. First, a finite sample of *N_s_* genomes is drawn from the population according to the genotype frequencies after selection and drift. Sequencing errors are then introduced by adding spurious mutations, modeled as a Poisson process with per-base error rate *err_seq_*. These errors are distributed across mutation classes according to the same categorization scheme used in the evolutionary model. The resulting observed frequencies therefore reflect both biological variation and technical noise.

#### Experimental parameter values

Model parameters were set to reflect the experimental design. The population size *N* = 2 ⋅ 10^7^ corresponds to the number of infecting virions per passage (Caspi et al., 2023), and sequencing coverage *N*_s_ was set according to the empirical data (*N*_s_ = 1309; see above). The probability that a mutation is synonymous was fixed based on the MS2 reference genome, and mutations were distributed across genes according to their coding lengths. The sequencing error rate *err_seq_* = 5 ⋅ 10^-5^ was set based on the reported accuracy of the long-read sequencing protocol (Callahan et al., 2021). All remaining evolutionary parameters, including mutation rate and fitness effects, were inferred from the data.

#### Two-stage inference strategy

A major challenge is that weakened purifying selection under high MOI could reflect either intrinsically mild deleterious effects or masking due to complementation. To separate these explanations, we adopted a two-stage inference strategy. In the first stage, the model was fitted to low-MOI data, where coinfection and complementation are assumed to be negligible. Gene-specific recessive probabilities were therefore fixed to zero, allowing inference of the intrinsic fitness effects of deleterious nonsynonymous mutations in each protein together with the baseline mutation and selection parameters. This stage yields full posterior distributions over these parameters.

In the second stage, the model was fitted to the high-MOI data. To reduce non-identifiability between intrinsic fitness effects and complementation, the protein-specific deleterious fitness parameters inferred in the first stage were fixed to their MAP estimates. For the remaining parameters, priors were informed by the first-stage posteriors, in particular, several global parameters (e.g., mutation rate and synonymous fitness effects) were assigned narrow uniform priors centered around their low-MOI MAP estimates, while other parameters retained broader priors. Inference in this stage therefore focused on the gene-specific probabilities that deleterious nonsynonymous mutations behave as recessive under coinfection, alongside the remaining free parameters. In this context, the recessive probability serves as a quantitative proxy for how shareable the corresponding protein function is within coinfected cells.

#### Simulation-based inference and model checking

Parameter inference was performed using neural posterior estimation (Greenberg et al., 2019) as implemented in the sbi framework (Tejero-Cantero et al., 2020). A masked autoregressive flow (MAF) (Papamakarios et al., 2017) was used as the neural density estimator. Summary statistics were based on the mean numbers of adaptive and non-adaptive nonsynonymous mutations per genotype for each protein, together with genome-wide adaptive and non-adaptive synonymous mutation counts. Finally, a single posterior distribution, conditioned on all three replicates, was inferred using the *Collective Posterior* (Ben Nun et al., 2026).

Posterior distributions were summarized using maximum-a-posteriori estimates and 95% highest density intervals, calculated by the ArviZ Python package (Kumar et al., 2019). To evaluate performance, the two inference stages were first tested on synthetic datasets generated under known parameter values (Prangle et al., 2014). We additionally performed posterior predictive checks by simulating data under inferred parameters and comparing simulated summary statistics with the observed passage-10 data. Finally, to test whether the coinfection model spuriously infers complementation in the absence of coinfection, we applied the second-stage framework to held-out low-MOI replicates in a leave-one-out control analysis.

## Code Availability

The source code is available at https://github.com/Stern-Lab/MS2_PG

## Acknowledgements

We thank Alison Feder and Uri Gophna for stimulating conversations that initiated this study. This study was supported by an ERC starting grant 852223 (RNAVirFitness) to A.S. This study was also supported by a fellowship to Y.M. and N.B.N. from the Edmond J. Safra Center for Bioinformatics at Tel Aviv University.

## Supplementary materials

**Figure S1.**
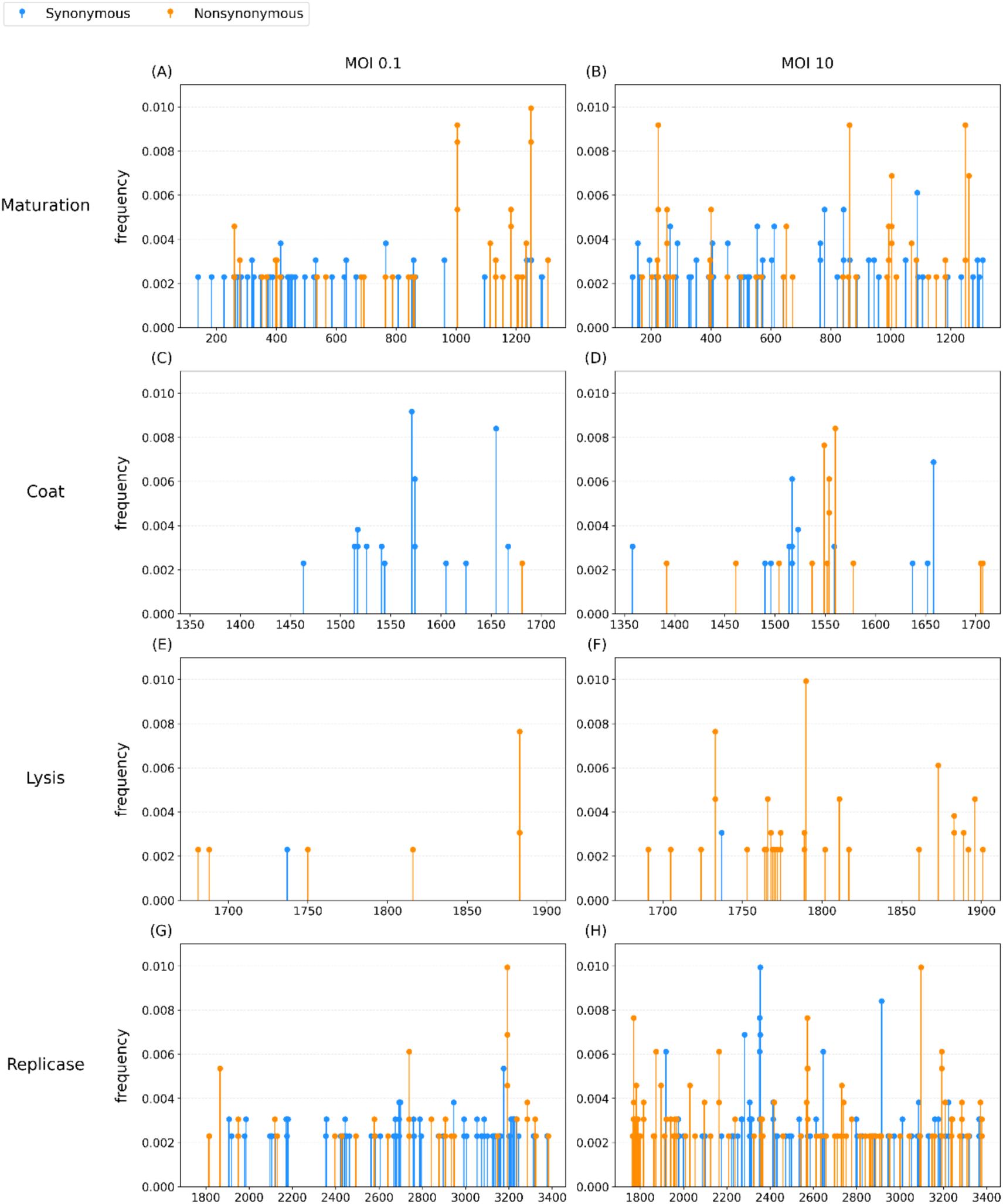
Low frequency mutation distributions across genes at low and high MOIs. Genomic positions of low-frequency mutations (frequency < 0.01) are shown for each gene under low (MOI 0.1, left column) and high (MOI 10, right column) infection conditions. Panels correspond to the four MS2 genes: maturation protein (Maturation; A-B), capsid protein (Coat; C-D), lysis protein (Lysis; E-F), and replicase (Replicase; G-H). Each dot represents a mutation observed in at least one independently evolved line, plotted by its genomic position and frequency. Synonymous mutations are shown in blue and nonsynonymous mutations in orange. For each gene, paired panels highlight differences in the spatial distribution and composition of low-frequency mutations between low and high MOI conditions.

**Figure S2.**
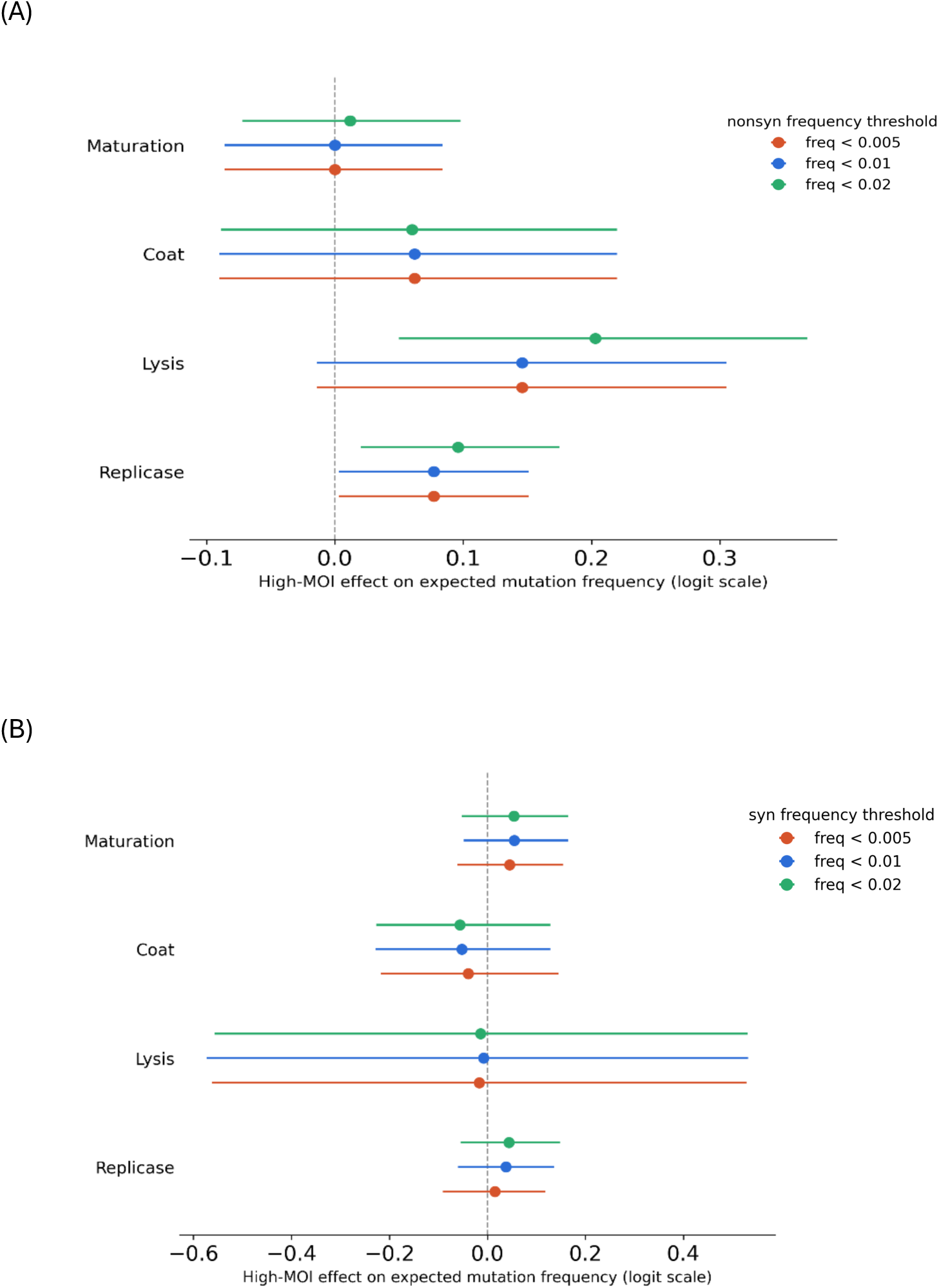
Sensitivity of hierarchical beta-binomial estimates to the low-frequency mutation threshold. Protein-specific high-MOI effects were estimated using hierarchical beta-binomial models fitted separately under three upper frequency thresholds: mutations below 0.5%, 1%, or 2% frequency. Effects are shown for (A) non-adaptive nonsynonymous mutations and (B) synonymous mutations. Points indicate posterior means and horizontal bars indicate 95% highest density intervals (HDIs). Effects are shown on the logit scale, where positive values indicate higher expected mutant-read frequencies under MOI = 10 than under MOI = 0.1. The dashed vertical line marks no high-MOI effect.

**Figure S3.**
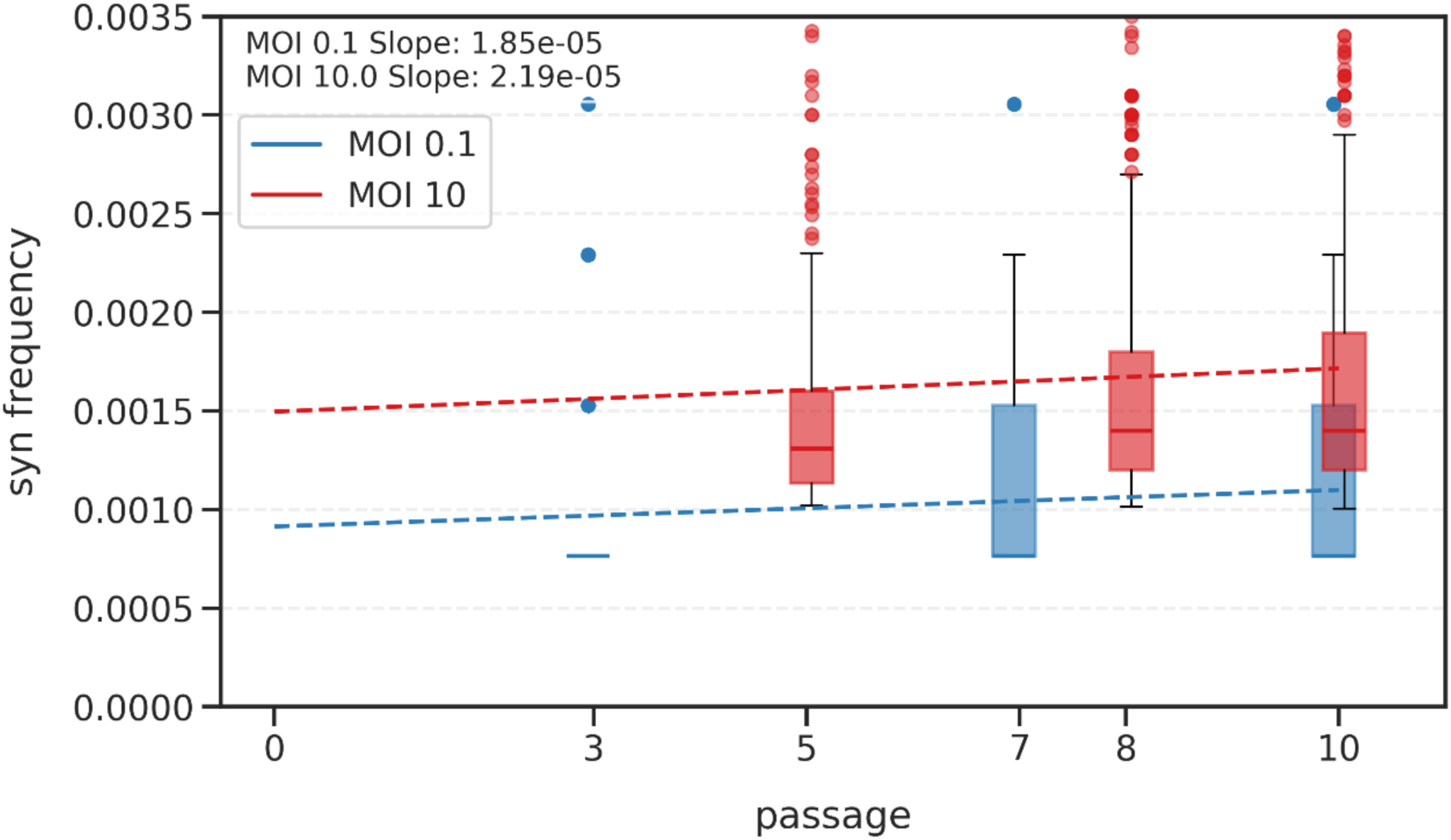
Accumulation of low-frequency synonymous mutations across passages. The accumulation of low-frequency synonymous mutations (f < 0.01) in the replicase and maturation proteins across serial passages of all three replicates under low and high MOIs. Linear regression was used to estimate the rate of synonymous mutation accumulation across passages, with dashed lines indicating the fitted slopes for each MOI condition. For the high MOI condition, mutation frequencies were calculated from short-read sequencing data, as long-read sequencing was available only for passage 10. The slight difference between the slopes suggests that MOI may influence the effective mutation accumulation rate, potentially reflecting differences in mutation rate and/or the number of replication cycles during infection.

**Figure S4.**
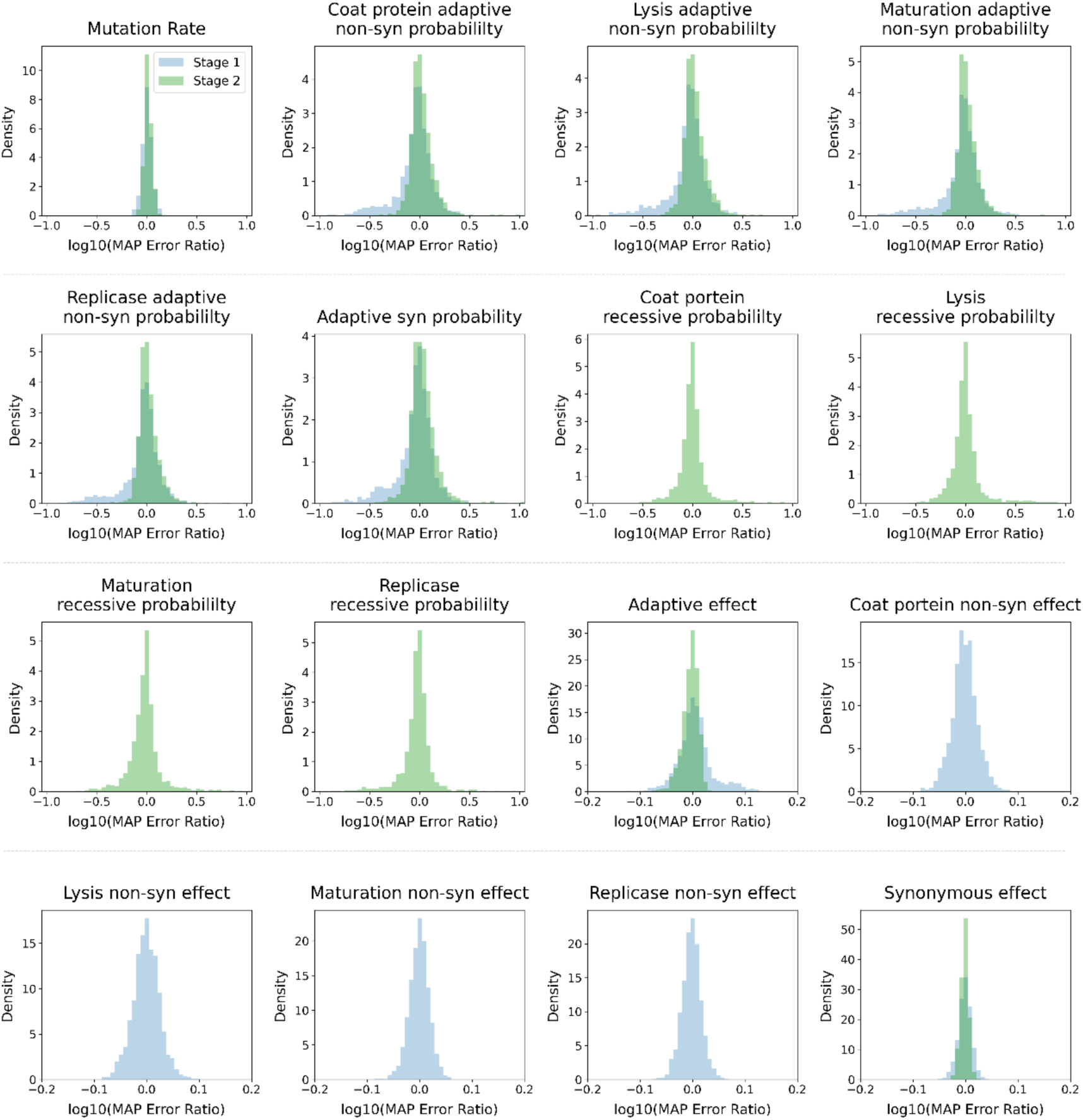
Synthetic-data benchmarks: MAP error ratios. Distributions of the log_10_ error ratio between MAP estimates and true parameter values across 2,000 simulations generated by the corresponding model. Results are shown for low MOI model (blue) and high MOI model (green), with overlapping distributions indicating parameters shared by both models. All distributions are mostly concentrated around zero which suggests the density estimators are not biased.

**Figure S5.**
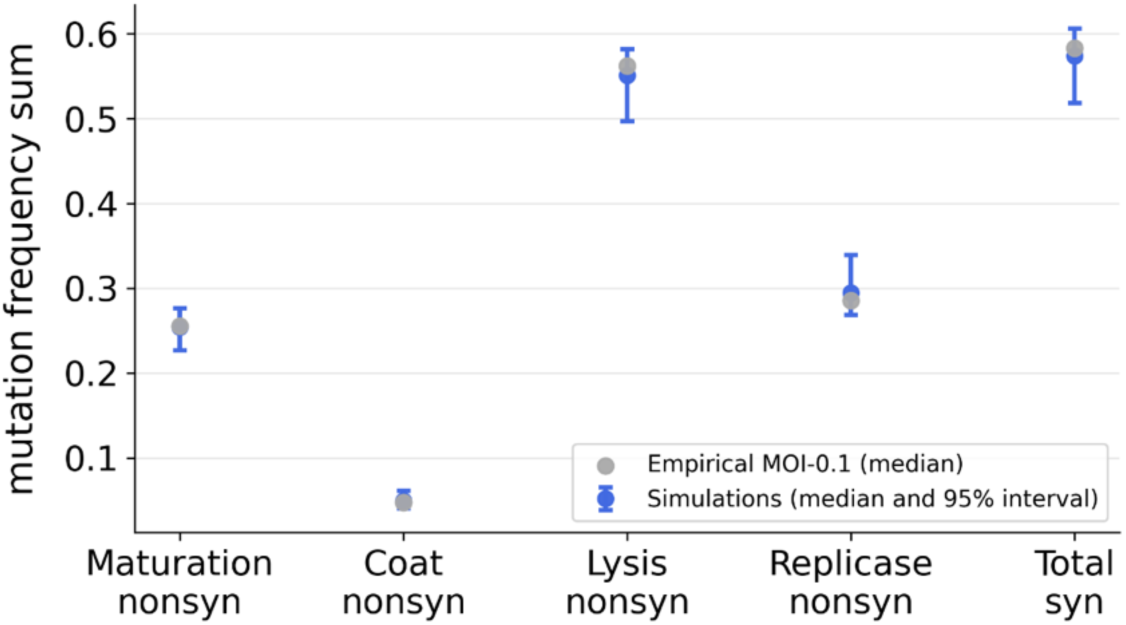
Posterior predictive checks of the inferred low-MOI evolutionary model. Comparison of empirical mutation frequencies sum at passage 10 (gray) to 200 simulations generated using parameters sampled from the inferred posteriors (blue) under MOI 0.1 across all mutation classes. Points show the median value for each mutation class, and blue error bars indicate the 95% simulation interval. For visualization, the ten-entry summary statistic was collapsed into five entries by summing adaptive and non-adaptive nonsynonymous frequencies within each protein, and by summing adaptive and non-adaptive synonymous frequencies genome-wide.

**Table S1.**
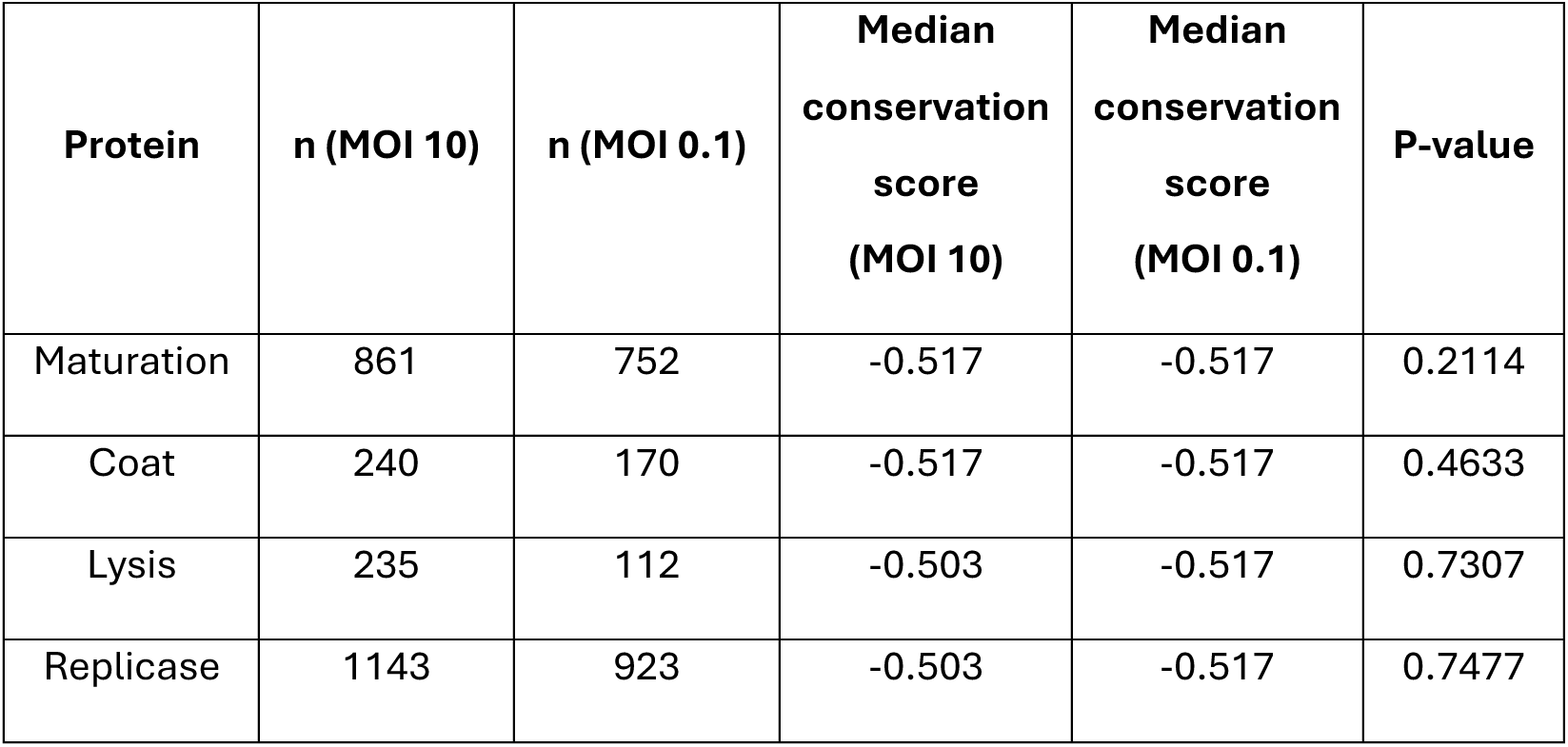
Comparison of conservation scores of mutated sites between MOI regimes. Site-specific conservation scores of nonsynonymous mutations observed under MOI 10 and MOI 0.1 were compared for each MS2 protein using a one-sided Mann-Whitney U test testing whether mutations under high MOI occur at more conserved sites (lower scores). Sample sizes (n) and median conservation scores are shown. No significant differences were detected for any protein (p > 0.05).

**Table S2.** Observed coverage on synthetic data. Coverage values report the proportion of cases the true parameter value fell within the nominal 95% HDI of the inferred posterior distribution. Coverage was evaluated using 2,000 synthetic datasets generated from each evolutionary model (MOI 0.1 and MOI 10), with each density estimator tested on data simulated from its corresponding model. Across parameters, coverage values are close to the nominal level of 0.95, though the estimators exhibit slight overconfidence, with empirical coverage consistently below 95%.

| parameter | Low MOI (0.1) model - coverage | High MOI (10) model - coverage |
| --- | --- | --- |
| $\mu$ | 0.9115 | 0.926 |
| $\omega_{syn}$ | 0.917 | 0.9165 |
| $\omega_{ada}$ | 0.937 | 0.9195 |
| $\omega_{ns}^{(mat)}$ | 0.921 | NA |
| $\omega_{ns}^{(cp)}$ | 0.93 | NA |
| $\omega_{ns}^{(lys)}$ | 0.93 | NA |
| $\omega_{ns}^{(rep)}$ | 0.9235 | NA |
| $p_{ns,ada}^{(mat)}$ | 0.9325 | 0.933 |
| $p_{ns,ada}^{(cp)}$ | 0.9375 | 0.9345 |
| $p_{ns,ada}^{(lys)}$ | 0.943 | 0.9245 |
| $p_{ns,ada}^{(rep)}$ | 0.9325 | 0.929 |
| $p_{syn,ada}$ | 0.9445 | 0.9285 |
| $p_{ns,rec}^{(mat)}$ | NA | 0.922 |
| $p_{ns,rec}^{(cp)}$ | NA | 0.9325 |
| $p_{ns,rec}^{(lys)}$ | NA | 0.9395 |
| $p_{ns,rec}^{(rep)}$ | NA | 0.9205 |

